# NCBP2-AS2 is a mitochondrial microprotein, regulates energy metabolism and neurogenesis, and is downregulated in Alzheimer’s disease

**DOI:** 10.1101/2025.01.25.634884

**Authors:** Stanislava Popova, Prabesh Bhattarai, Elanur Yilmaz, Daniela Lascu, Juo-Han Kuo, Gizem Erdem, Basak Coban, Jitka Michling, Mehmet Ilyas Cosacak, Huseyin Tayran, Thomas Kurth, Alexandra Schambony, Frank Buchholz, Marc Gentzel, Caghan Kizil

## Abstract

Microproteins, short functional peptides encoded by small genes, are emerging as critical regulators of cellular processes, yet their roles in mitochondrial function and neurodegeneration remain underexplored. In this study, we identify NCBP2-AS2 as an evolutionarily conserved mitochondrial microprotein with significant roles in energy metabolism and neurogenesis. Using a combination of cellular and molecular approaches, including CRISPR/Cas9 knockout models, stoichiometric co- immunoprecipitation, and advanced imaging techniques, we demonstrate that NCBP2-AS2 localizes to the inner mitochondrial space and interacts with translocase of the inner membrane (TIM) chaperones. These interactions suggest a role in ATPase subunit transport, supported by the observed reductions in ATPase subunit levels and impaired glucose metabolism in NCBP2-AS2-deficient cells. In zebrafish, NCBP2-AS2 knockout led to increased astroglial proliferation, microglial abundance, and enhanced neurogenesis, particularly under amyloid pathology. Notably, we show that NCBP2-AS2 expression is consistently downregulated in human Alzheimer’s disease brains and zebrafish amyloidosis models, suggesting a conserved role in neurodegenerative pathology. These findings reveal a novel link between mitochondrial protein transport, energy metabolism, and neural regeneration, positioning NCBP2-AS2 as a potential therapeutic target for mitigating mitochondrial dysfunction and promoting neurogenesis in neurodegenerative diseases such as Alzheimer’s disease.

## Introduction

Microproteins are small, functional peptides encoded by short genes that play key roles in development, signaling, and disease (Hassel et al., 2023, Staudt and Wenkel, 2011, Wu et al., 2022). Advances in genome sequencing and bioinformatics have enabled identification and study of these small proteins, which are often overlooked due to their small size and encoding by short open reading frames (sORFs). Techniques such as ribosome profiling and transcriptomics have been instrumental in identifying sORFs that are actively translated into functional proteins, revealing a previously underexplored area of the proteome (Hassel et al., 2023). Building on the emerging understanding of microproteins, mitochondrial microproteins have gained significant attention for their roles in regulating energy metabolism, mitochondrial respiration, and cell signalling (Anderson et al., 2015, Makarewich et al., 2018b, Rodrigues et al., 2021, Makarewich et al., 2018a, Wang et al., 2020, Lin et al., 2019, Stein et al., 2018, Zhang et al., 2020, Miller et al., 2022, Zhao et al., 2020, Benezra et al., 1990, Ikeda et al., 2013). For instance, Mitoregulin and Brawnin influence respiratory efficiency by mediating the assembly of mitochondrial complexes (Stein et al., 2018, Zhang et al., 2020), while SHMOOSE has been implicated in mitochondrial respiration and its potential role in Alzheimer’s disease (AD) (Miller et al., 2022). These examples underscore the importance of microproteins in mitochondrial function and their potential contributions to human diseases. A particular relevance of mitochondrial dysfunction to disease is observed in neurodegenerative diseases, including AD. Recent studies have highlighted the significance of mitochondrial proteins in regulating neuronal health and degeneration has been linked to mitochondrial dysfunction and respiration defects in AD, suggesting that mitochondrial microproteins may contribute to the pathophysiology of this disease (Mor et al., 2020, Liu et al., 2015, Gray et al., 2014, Askenazi et al., 2023, Miller et al., 2022, Kalia et al., 2022, March-Diaz et al., 2021, Bai et al., 2021, Zhang et al., 2015, Kawatani et al., 2023).

The long non-coding RNA (lncRNA) NCBP2-AS2 has been shown to encode a microprotein implicated in diverse biological processes, including its potential role as a mitochondrial microprotein contributing to energy metabolism and cellular signaling. Previous studies have demonstrated its involvement in tumorigenesis and angiogenesis (Kugeratski et al., 2019, Xu et al., 2020). For example, the microprotein HIAR, encoded by NCBP2-AS2, was identified as a hypoxia-induced angiogenesis regulator in mammary cancer (Kugeratski et al., 2019), while KRASIM, another microprotein derived from NCBP2-AS2, inhibits ERK signaling activity in hepatocellular carcinoma (Xu et al., 2020). However, its role in mitochondria and its relevance to neurodegenerative diseases have remained unexplored. In this study, we provide a detailed characterization of NCBP2-AS2 as a mitochondrial microprotein, to explore its novel roles in mitochondrial localization, energy metabolism, and neurodegeneration. Using a combination of immunocytochemistry, electron microscopy, and CRISPR/Cas9 knockout models, we demonstrate that NCBP2-AS2 localizes to mitochondria and interacts with mitochondrial translocase complexes. Loss of NCBP2-AS2 function alters mitochondrial energy metabolism, specifically reducing ATPase subunit levels and glucose utilization. Furthermore, NCBP2-AS2 knockout in zebrafish reveals its role in regulating astroglial proliferation, microglial abundance, and neurogenesis, with a significant increase in neurogenic outcomes observed in the knockout model. Notably, we also show that NCBP2-AS2 expression is downregulated in AD patients and animal models, highlighting its potential role in neurodegeneration. Overall, our findings establish NCBP2-AS2 as a conserved mitochondrial microprotein with critical functions in energy metabolism and neurogenesis. These results suggest that mitochondrial microproteins like NCBP2-AS2 may represent novel therapeutic targets for addressing mitochondrial dysfunction in neurodegenerative diseases such as AD, and future research could explore its potential for developing targeted therapies or its broader role in other mitochondrial-related diseases.

## Results

### NCBP2-AS2 is a mitochondrial protein

To determine the cellular localization of NCBP2-AS2, we employed a bacterial artificial chromosome (BAC) tagging approach to generate a reporter HeLa cell line expressing the GFP-tagged NCBP2-AS2 open reading frame in its genomic context (**Figure 1A, Data S1**). Western blot analysis confirmed the expression of NCBP2-AS2-GFP in the HeLa BAC line, with a protein band corresponding to the expected molecular weight, whereas no such band was detected in wild-type HeLa cells (**Figure 1B**). GAPDH served as a loading control. Live-cell imaging of the BAC-modified HeLa cells revealed GFP fluorescence predominantly in a perinuclear pattern consistent with mitochondrial localization (**Figure 1C**). Immunocytochemistry on fixed cells showed that NCBP2-AS2-GFP co-localized with TOMM20, a mitochondrial outer membrane marker (**Figure 1D**). Mitochondrial localization was further confirmed by subcellular fractionation, where NCBP2-AS2-GFP was enriched in the mitochondrial fraction alongside PAM16, a known mitochondrial marker, with minimal presence in lysate and supernatant fractions (**Figure 1E**). To validate the mitochondrial localization at ultrastructural resolution, we performed transmission electron microscopy (TEM) coupled with GFP immunogold labeling. Immunogold particles marking NCBP2-AS2-GFP were localized to mitochondria, confirming its mitochondrial localization (**Figure 1F**). Higher magnification of the mitochondria further highlighted the specific distribution of NCBP2-AS2-GFP within these organelles (**Figure 1F**’). These results demonstrate that NCBP2-AS2 is a mitochondrial protein localized to this organelle.

**Figure 1:**
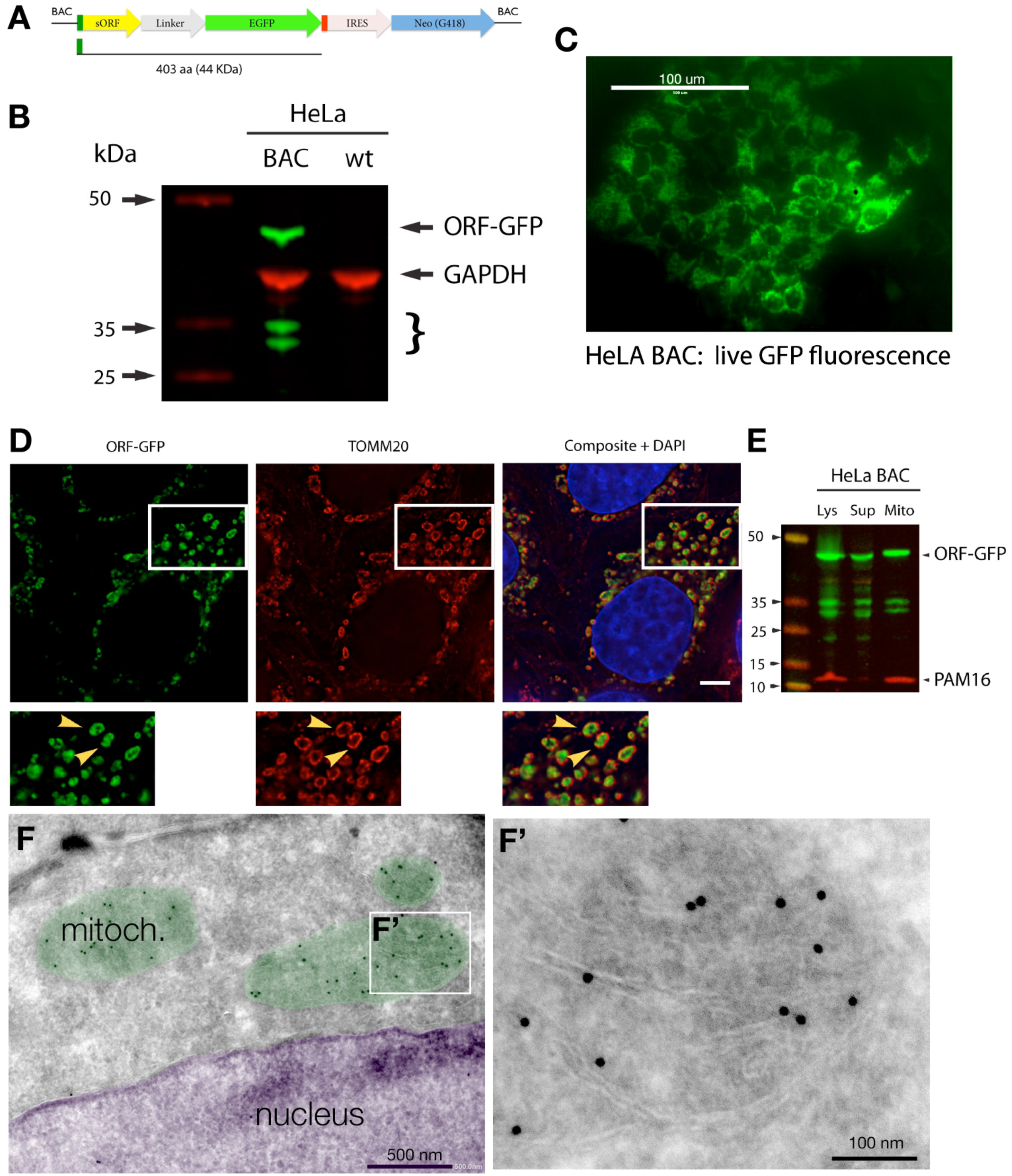
Validation and localization of NCBP2-AS2-GFP expressed in HeLa cells. (A) Schematic representation of the BAC recombineering construct, showing the open reading frame (ORF) of NCBP2- AS2 fused to GFP (green), followed by an IRES sequence and a neomycin resistance cassette (Neo; blue). The ORF encodes a 44 kDa fusion protein. (B) Western blot analysis of HeLa cells expressing the BAC construct (BAC) compared to wild-type cells (wt). The NCBP2-AS2-GFP fusion protein is detected in green, while GAPDH serves as a loading control (red). (C) Live fluorescence microscopy of HeLa BAC cells reveals perinuclear localization of GFP-tagged NCBP2-AS2. Scale bar: 100 μm. (D) Immunocytochemistry shows colocalization of the NCBP2-AS2-GFP fusion protein (green) with the mitochondrial marker TOMM20 (red). Nuclei are counterstained with DAPI (blue). Insets highlight higher magnification of colocalized regions, indicated by yellow arrowheads. Scale bar: 5 μm. (E) Subcellular fractionation confirms enrichment of NCBP2-AS2-GFP in the mitochondrial fraction, as shown by co- fractionation with the mitochondrial marker PAM16. (F) Transmission electron microscopy (TEM) with immunogold labeling of cryo-sections identifies NCBP2-AS2-GFP localized within mitochondria (mitoch.). (F’) Higher magnification of the boxed region in (F) confirms the mitochondrial localization of the fusion protein. Scale bars: 500 nm in (F) and 100 nm in (F’).

### NCBP2-AS2 interacts with mitochondrial inner membrane transport complexes and regulates energy metabolism

To identify the interaction partners and functional role of NCBP2-AS2, we generated CRISPR/Cas9- based knockout (KO) lines in HEK293T cells. Three clones with consistent genomic deletions were created, resulting in significant reduction of endogenous NCBP2-AS2 expression compared to wild-type cells (**Figure 2A, B; Data S1, Data S2**). Clone 2 was chosen for subsequent experiments due to its consistent genomic deletion and significant reduction in NCBP2-AS2 expression. To investigate NCBP2-AS2 interaction partners, we expressed FLAG-tagged NCBP2-AS2 alongside a control GFP-expressing plasmid in stoichiometrically varying transfection ratios (**Figure 2C**). By doing so, different levels of NCBP2-AS2 would be expressed in the cells and true co-immunoprecipitated interaction partners would also show a parallel stoichiometry in the resulting mass spectrometric analyses. Western blot analysis of immunoprecipitated samples confirmed successful dosing of NCBP2-AS2, as lower levels of NCBP2- AS2 in the input fraction correlated with reduced elution from the beads and decreased supernatant levels (**Figure 2D**).

**Figure 2:**
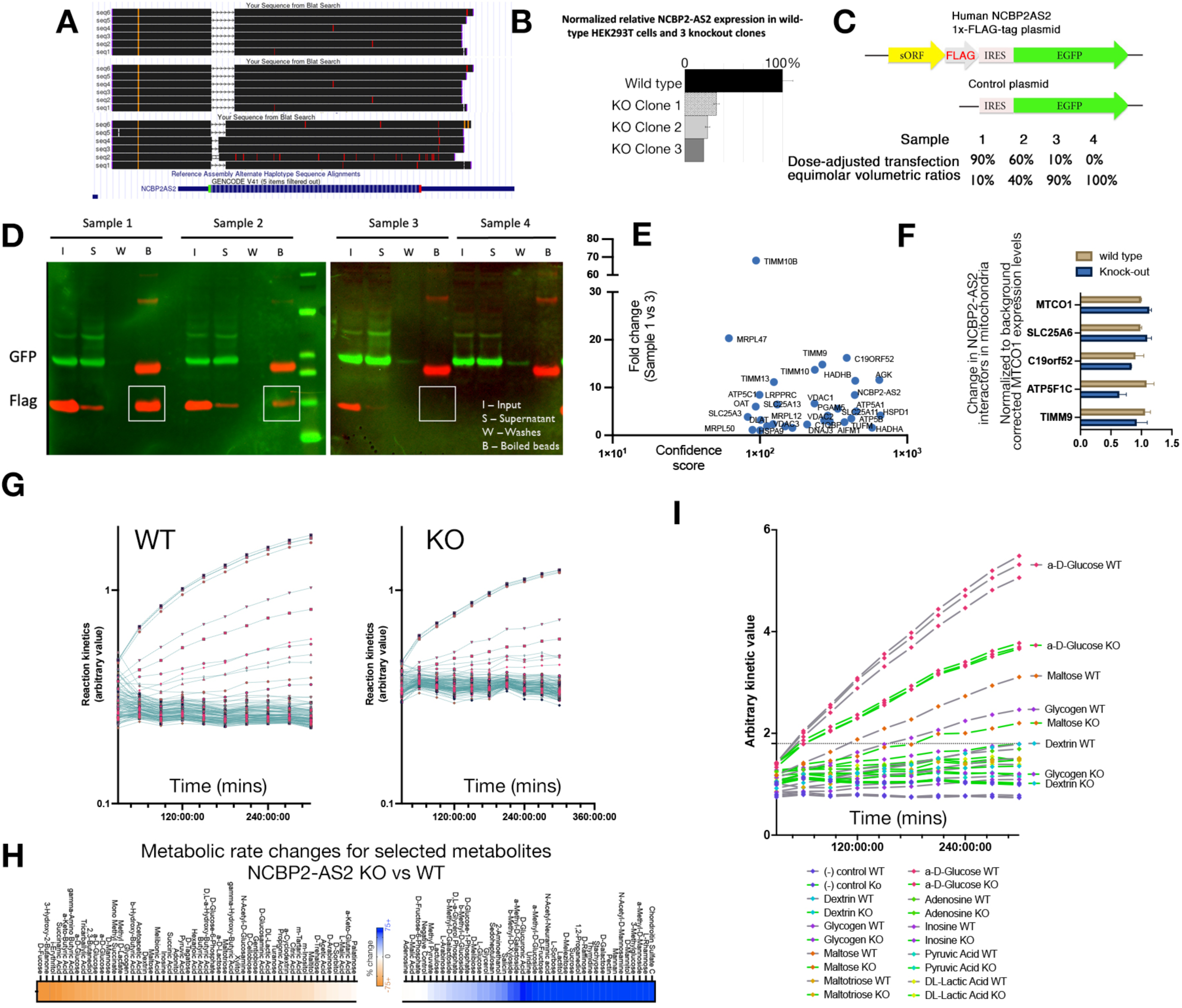
Functional analysis of NCBP2-AS2 deletion in HEK293T cells. (A) Sequencing results for three CRISPR/Cas9-generated NCBP2-AS2 knockout (KO) clones in HEK293T cells. Six replicate sequencing reads were performed for each clone, confirming successful deletion. (B) Relative RNA expression levels of NCBP2-AS2 in wild-type (WT) and KO clones. Expression was normalized to wild-type levels (100%), demonstrating complete knockout in all clones. (C) Schematic of the stoichiometry-adjusted co-immunoprecipitation (co-IP) strategy, highlighting FLAG- and GFP-tagged NCBP2-AS2 constructs and dose-adjusted plasmid transfection ratios for samples 1–4. (D) Representative western blots of co-IP experiments. GFP- and FLAG-tagged NCBP2-AS2 constructs were analyzed in input (I), supernatant (S), wash fractions (W), and boiled bead eluates (B). White boxes indicate regions containing GFP- or FLAG-tagged NCBP2-AS2. (E) Identification of mitochondrial co-IP interaction partners of NCBP2-AS2. Proteins are plotted by fold change (Sample 1 vs. 3) and confidence score, with key interactors labeled. (F) Quantification of changes in the abundance of protein products from select mitochondrial interaction partners in WT vs. KO cells. (G) Time-course substrate utilization kinetics for WT and KO cells, showing relative utilization rates across a 6-hour time frame. Data from all tested substrates are shown. (H) Heatmap showing metabolic rate changes for selected metabolites in KO versus WT cells. Metabolites involved in carbohydrate metabolism are prominently affected. (I) Specific substrate kinetic comparisons between WT and KO clones, focusing on glucose, maltose, glycogen, dextrin, and other key metabolites.

LC-MS/MS analysis of co-immunoprecipitated proteins revealed 31 high-confidence interaction partners of NCBP2-AS2, primarily mitochondrial proteins, including self-interactions (**Figure 2E, Data S3**). Among the top interactors were components of the translocase of the inner mitochondrial membrane (TIM) complex, such as TIMM9, TIMM10, and TIMM10B. Additional interactors included C19ORF52 (TIMM29), which bridges the inner and outer mitochondrial membranes (Kang et al., 2016), as well as TIMM8 and TIMM13, which are involved in multi-pass transmembrane protein translocation (Paschen et al., 2000, Beverly et al., 2008). Proteins from oxidative phosphorylation complexes were also identified, including ATP5A1, ATP5B, and ATP5F1C (Complex V ATPase subunits), along with all three voltage-dependent anion channels (VDAC1, VDAC2, and VDAC3). These interactions suggest that NCBP2-AS2 is involved in mitochondrial protein translocation and ATPase subunit transport.

To test whether NCBP2-AS2 regulates mitochondrial protein levels, we performed mitochondrial fractionation and western blot analyses of selected candidate proteins in wild-type and KO HEK293T cells. Loss of NCBP2-AS2 resulted in a significant reduction of ATP5C1 subunit expression in the mitochondrial fraction, normalized to MTOC1 as a loading control (**Figure 2F; Data S4**). Given the reduction of ATP5C1, we hypothesized that NCBP2-AS2 knockout would alter glucose metabolism. Metabolic assays revealed significant changes in the utilization of glucose and related substrates (glycogen, maltose, and dextrin) in NCBP2-AS2 KO cells compared to wild-type cells (**Figure 2G-I; Data S5**). Kinetic analyses demonstrated reduced glucose utilization in KO cells, indicating impaired energy metabolism. These results suggest that NCBP2-AS2 interacts with mitochondrial translocase complexes and may regulate the transport and function of ATPase subunits, thereby influencing glucose metabolism and energy production.

### NCBP2-AS2 is evolutionarily conservation at its localization to mitochondria

To investigate the evolutionary conservation and mitochondrial localization of NCBP2-AS2, we aligned zebrafish, human, and mouse NCBP2-AS2 protein sequences using Clustal-W. The alignment revealed highly conserved stretches of amino acid sequences, particularly within regions predicted to influence mitochondrial targeting (**Figure 3A**). In silico localization prediction algorithms supported these findings, predicting zebrafish NCBP2-AS2 localization to mitochondria via mitochondrial targeting signals (**Figure 3B**). To experimentally validate the localization, expression plasmids containing human or zebrafish NCBP2-AS2 fused with a FLAG tag were generated (**Data S1**). Transfection of these constructs into HEK293T cells showed co-localization of both human and zebrafish NCBP2-AS2 with mitochondrial markers TOMM20 and PAM16, as visualized by immunocytochemistry (**Figure 3C**). Control plasmid-transfected and non-transfected cells exhibited no such mitochondrial localization, confirming the specificity of the tagged NCBP2-AS2 localization. Merged images and higher magnification insets further demonstrated clear overlap of the NCBP2-AS2 signal with mitochondrial markers. We next assessed the expression of NCBP2-AS2 in the adult zebrafish brain by analyzing previously published single-cell transcriptomics datasets. NCBP2-AS2 expression was detected in neurons (marked by *sv2a*) and astroglia (marked by *s100b*), confirming its presence in both cell types critical to brain function (**Figure 3D**). PCR analysis of zebrafish brain RNA validated the expression of NCBP2-AS2 alongside control genes *eef1a1* and *actb2* (**Figure 3E**).

**Figure 3:**
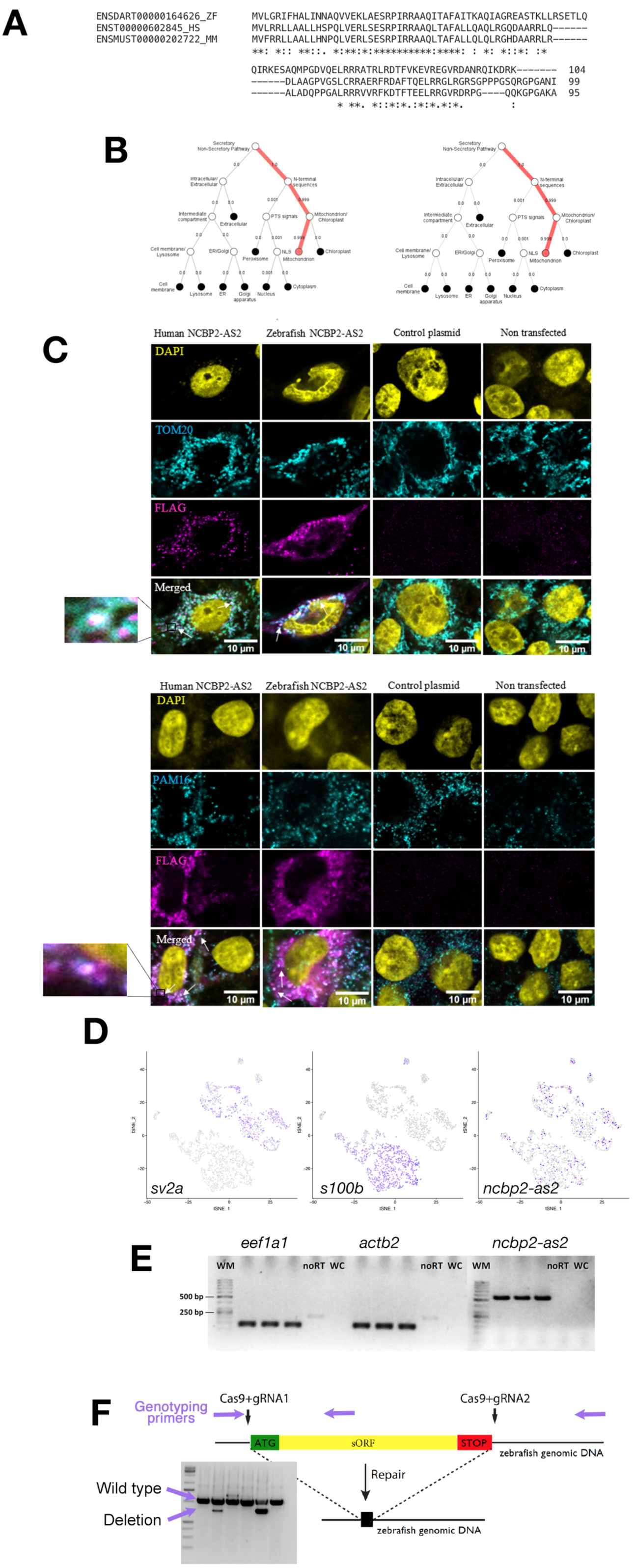
Comparative and functional characterization of NCBP2-AS2 in zebrafish, mouse, and humans. (A) Clustal-W multiple sequence alignment of NCBP2-AS2 protein orthologs in zebrafish, mouse, and humans. Highly conserved regions are indicated by asterisks, showcasing sequence similarity across species. (B) In silico localization analysis predicts zebrafish NCBP2-AS2 protein to be mitochondrial. Pathways involving mitochondrial targeting signals are highlighted in red. (C) Co- localization of FLAG-tagged human and zebrafish NCBP2-AS2 with mitochondrial markers TOMM20 and PAM16 in HEK293T cells. DAPI staining indicates nuclei (yellow). Insets show higher magnification of co-localized regions (arrows). Scale bars: 10 μm. (D) Single-cell tSNE plots illustrating expression patterns of neuronal marker *sv2a*, glial marker *s100b*, and *ncbp2-as2* . NCBP2-AS2 is expressed in both neurons and glial cells in zebrafish. (E) RT-PCR analysis confirms expression of NCBP2-AS2 in zebrafish brain tissue. Amplification of control genes eef1a1 and actb2 is shown alongside NCBP2-AS2. WM: weight marker, noRT: no reverse transcription control, WC: water control. (F) CRISPR-Cas9 strategy for deletion of NCBP2-AS2 in zebrafish. The schematic shows Cas9 cleavage sites flanking the ORF. PCR genotyping primers (purple arrows) detect wild-type (WT), heterozygous deletion, and homozygous knockout genotypes. Representative gel electrophoresis confirms the genotypes.

### Zebrafish knockout for NCBP2-AS2 alters astroglial proliferation and neurogenesis in health and under Alzheimer’s disease pathology

To explore the functional role of NCBP2-AS2 *in vivo*, we generated a CRISPR/Cas9-mediated knockout of the NCBP2-AS2 locus in zebrafish. Two guide RNAs (gRNAs) flanking the open reading frame (ORF) were used to induce deletions. PCR genotyping confirmed successful generation of homozygous deletion lines, as shown by the absence of the wild-type band and the presence of a deletion-specific band (**Figure 3F**).

To analyze the role of NCBP2-AS2 in the brain, we examined the effects of the knockout effects on astroglial proliferation, microglial abundance, cell death, and neurogenesis in wild-type (WT) and NCBP2-AS2 knockout (KO) zebrafish at 6 and 11 months post-fertilization (mpf). Immunohistochemistry showed that BrdU-positive S100β+ astroglia, indicative of proliferating astrocytes, were significantly increased in KO zebrafish at 6 mpf compared to WT (**Figure 4A; Data S6**). At 11 mpf, this difference was no longer observed, suggesting and consistent with age-dependent effects on astroglial proliferation (Bhattarai et al., 2017a, Bhattarai et al., 2017b). We also examined microglial abundance using L-Plastin staining and observed a significant increase in microglial numbers in KO zebrafish at both 6 and 11 mpf (**Figure 4B**). Concurrently, TUNEL staining revealed a significant rise in apoptotic nuclei in KO zebrafish at both time points, indicating increased cell death in the absence of NCBP2-AS2 (**Figure 4B**). To determine whether increased astroglial proliferation influences neurogenesis, we assessed the overlap between glial (S100β) and neuronal (HuC/D) markers, indicative of transitioning progenitors to pro-neural fate. Immunohistochemistry revealed a significant increase in the percentage of S100β+/HuC/D+ double-positive cells in KO zebrafish compared to WT at 6 mpf (**Figure 4C, D**). Additionally, BrdU labeling demonstrated a significant increase in BrdU+/HuC/D+ newborn neurons in KO zebrafish, suggesting enhanced neurogenic outcomes in the absence of NCBP2-AS2 (**Figure 4E, F**). These results indicate that NCBP2-AS2 knockout leads to increased astroglial proliferation, microglial abundance, and neurogenesis in zebrafish brains. The age-dependent effects on astroglial proliferation and persistent increase in neurogenic outcomes highlight the regulatory role of NCBP2-AS2 in brain homeostasis and cell turnover.

**Figure 4:**
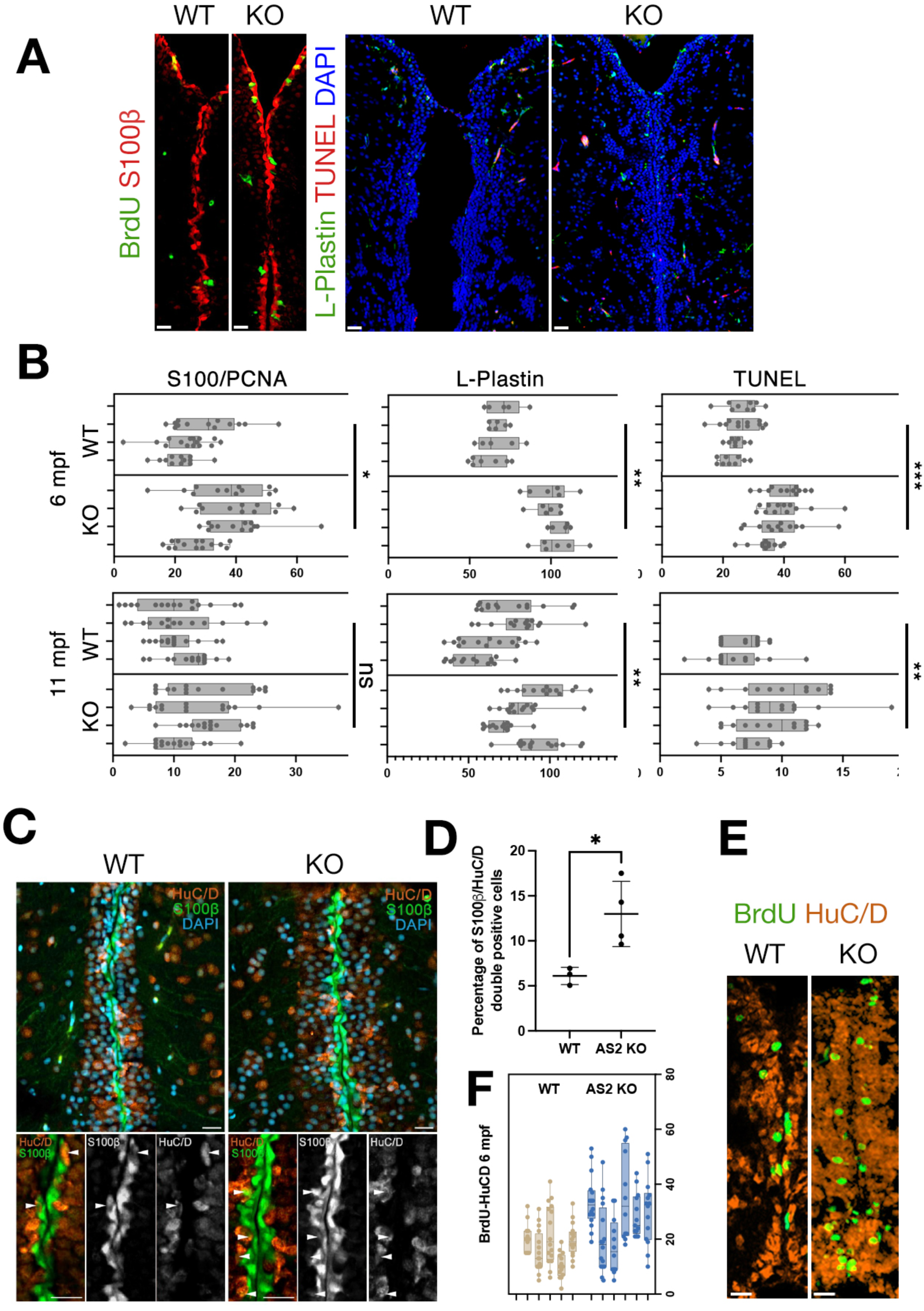
Analysis of glial proliferation, microglia activity, apoptosis, and neurogenesis in NCBP2-AS2 knockout zebrafish. (A) Immunohistochemical staining of zebrafish brains. Left panel: S100β (red; astroglia) and PCNA (green; proliferating cells). Right panel: L-Plastin (green; microglia) and TUNEL (red; apoptotic cells). DAPI marks nuclei (blue). Scale bars: 25 μm. (B) Quantification of S100β/PCNA-positive astroglia, L-Plastin-positive microglia, and TUNEL-positive apoptotic cells in 6- and 11-month-old zebrafish brains. Wild-type (WT) and NCBP2-AS2 knockout (KO) animals were compared. Significant differences are indicated (*p < 0.05, **p < 0.01, ***p < 0.001). (C) Immunohistochemical analysis showing HuC/D (orange; early neurons), S100β (green; astroglia), and DAPI-stained nuclei in WT and KO zebrafish brains. Insets highlight colocalization of S100β and HuC/D. White arrowheads indicate representative double-positive cells. (D) Quantification of S100β/PCNA double-positive cells in WT and KO zebrafish brains, showing a significant increase in astroglial proliferation upon NCBP2-AS2 knockout (*p < 0.05). (E) Immunostaining for BrdU (green; proliferating cells) and HuC/D (red; early neurons) shows enhanced neurogenesis in NCBP2-AS2 KO zebrafish compared to WT controls. Scale bars: 25 μm. (F) Quantification of BrdU+/HuC/D+ newborn neurons at 6 months post-fertilization (mpf) indicates increased neurogenesis in NCBP2-AS2 KO zebrafish.

### NCBP2-AS2 is downregulated in Alzheimer’s disease patients and zebrafish model of amyloidosis, and it regulates astroglial proliferation and neurogenesis in zebrafish amyloidosis model

Mitochondrial dysfunction is highly relevant to the pathogenesis of Alzheimer’s disease (AD), highlighting the relevance of mitochondrial proteins like NCBP2-AS2 in this neurodegenerative conditions (Kawatani et al., 2023, March-Diaz et al., 2021, Bai et al., 2021, Yang et al., 2015). To explore whether NCBP2-AS2 expression is altered in Alzheimer’s conditions, we utilized published single-cell sequencing (scSeq) and single-nucleus sequencing (snSeq) datasets from zebrafish (Tayran et al., 2024) and human brains (Is et al., 2024). These analyses revealed a consistent reduction in NCBP2-AS2 expression in both astrocytes and neurons from human AD patients and zebrafish amyloid models compared to controls (**Figure 5A- C**). This observation suggests that NCBP2-AS2 downregulation is a conserved feature across species.

**Figure 5.**
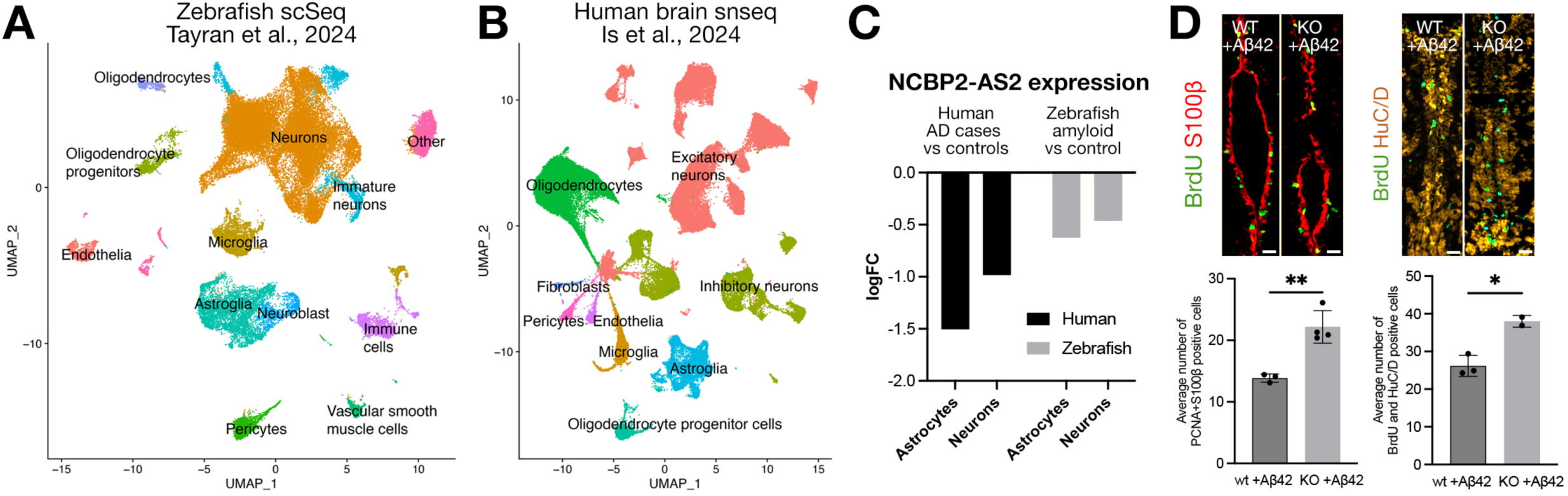
NCBP2-AS2 reduces in Alzheimer’s disease, and NCBP2-AS2 knockout increases proliferation and neurogenesis in amyloidosis model in zebrafish. (A) UMAP visualization of single- cell sequencing (scSeq) data from the zebrafish brain, showing clustering of various cell types, including neurons, oligodendrocytes, astroglia, and microglia. (B) UMAP plot of single-nucleus sequencing (snSeq) data from the human brain, displaying clusters of excitatory and inhibitory neurons, astroglia, microglia, and other cell types. (C) Expression levels of NCBP2-AS2 in astrocytes and neurons from human Alzheimer’s disease (AD) cases vs. controls (black bars) and zebrafish amyloid vs. control models (gray bars). The data are represented as log fold changes (logFC), showing a decrease in NCBP2-AS2 expression in both species. (D) Immunohistochemistry images (upper panels) showing BrdU-positive cells co-stained with S100β (left) or HuC/D (right) in wild-type (WT) and knockout (KO) zebrafish brains treated with Aβ42. Quantification (lower panels) reveals significantly increased numbers of BrdU-positive S100β+ (astrocytic marker) and HuC/D+ (neuronal marker) cells in KO+Aβ42 zebrafish compared to WT+Aβ42. Scale bars, 10 μm. Data are presented as mean ± SEM. Statistical significance: *p < 0.05, **p < 0.01.

To investigate the functional consequences of NCBP2-AS2 knockout (KO) in an amyloidosis context, we examined neurogenesis and astroglial proliferation in zebrafish brains treated with amyloid beta (Aβ42), a model that has remarkable similarities to amyloid pathology in humans (Kizil and Bhattarai, 2018, Kizil, 2018, Cosacak et al., 2019, Cosacak et al., 2020a, Cosacak et al., 2022) and has been used to functionally analyze AD-related genes and genetic variants in humans reliably (Tayran et al., 2024, Haage et al., 2024, Bhattarai et al., 2024, Is et al., 2024, Ray et al., 2023, Kim et al., 2023b, Kim et al., 2023a, Felsky et al., 2023, Kizil et al., 2022, Lee et al., 2022, Cosacak et al., 2022, Turgutalp et al., 2022, Bhattarai et al., 2022, Siddiqui et al., 2021, Turgutalp and Kizil, 2024). Immunohistochemical staining revealed a significant increase in BrdU-positive S100β+ astrocytes and BrdU-positive HuC/D+ neurons in KO+Aβ42 zebrafish compared to WT+Aβ42 zebrafish (**Figure 5D**). These findings suggest that the loss of NCBP2-AS2 enhances both astroglial proliferation in the presence of amyloid pathology. Furthermore, quantitative analysis demonstrated that the NCBP2-AS2 KO zebrafish increased neurogenesis, indicating a compensatory regenerative response to amyloidosis (**Figure 5D, E**). These data support the hypothesis that NCBP2-AS2 negatively regulates neurogenic processes, and its downregulation in AD may represent an adaptive mechanism to counteract neurodegenerative damage. The reduction of NCBP2-AS2 in AD and its impact on neurogenesis and astroglial dynamics in amyloidosis models underscore the protein’s dual role in mitochondrial function and brain resilience. These findings highlight the potential of NCBP2-AS2 as a therapeutic target for modulating neurogenesis and mitigating the effects of mitochondrial dysfunction in AD.

## Discussion

In this study, we establish NCBP2-AS2 as a mitochondrial microprotein with critical roles in energy metabolism and neurogenesis. We suggest that NCBP2-AS2 is a novel component of the mitochondrial inner membrane transport system, specifically interacting with translocase of the inner membrane (TIM) chaperones. Identifying NCBP2-AS2 as part of this system is significant because TIM chaperones play a critical role in maintaining mitochondrial protein homeostasis and facilitating the import and assembly of essential metabolic complexes, processes that are central to mitochondrial function and cellular energy metabolism (Grevel and Becker, 2020, Kang et al., 2016, Beverly et al., 2008, Xu et al., 2005, Paschen et al., 2000). Co-immunoprecipitation experiments demonstrated high-confidence interactions with TIM complex proteins such as TIMM9, TIMM10, and TIMM10B, as well as with components of oxidative phosphorylation complexes, including ATPase subunits. This implicates NCBP2-AS2 in protein translocation and ATPase subunit transport, supported by the reduction of ATP5C1 levels in knockout (KO) cells. Localization analyses via electron microscopy confirmed its presence in the inner mitochondrial space (IMS), consistent with its proposed role in mitochondrial transport dynamics. Although the interaction between NCBP2-AS2 and the TIM complex may reflect its own protein import, our results also suggest a functional role in ATPase subunit transport, as evidenced by metabolic alterations in glucose utilization in KO cells. Specifically, the reduced glucose metabolism observed in KO cells aligns with decreased levels of ATPase subunits, implicating NCBP2-AS2 in maintaining efficient mitochondrial energy production through proper subunit translocation. Further structural and functional studies are needed to delineate the exact mechanisms underlying these interactions.

In addition to mitochondrial interactions, we found NCBP2-AS2 to interact with nuclear and cytoskeletal proteins, including actin and kinesins. These interactions suggest that NCBP2-AS2 may influence broader cellular processes such as mitochondrial dynamics, intracellular transport, and cellular localization. Actin and kinesins are key regulators of cytoskeletal architecture and intracellular cargo transport, which are essential for maintaining mitochondrial function and subcellular positioning (Kim and Cheong, 2020). Exploring how these interactions contribute to NCBP2-AS2’s roles beyond mitochondrial function could reveal additional layers of its physiological significance. These proteins are known to regulate mitochondrial dynamics and subcellular localization (Kim and Cheong, 2020). While these interactions may indicate broader cellular roles, electron microscopy did not detect NCBP2-AS2 in the nucleus, suggesting that such interactions may be transient. Future studies should explore the stability and functional relevance of these interactions to clarify their contribution to NCBP2-AS2’s physiological roles.

NCBP2-AS2 had an impact on astroglial proliferation and neurogenesis in zebrafish KO models. Increased astroglial proliferation at 6 months post-fertilization (mpf) was accompanied by enhanced neurogenic outcomes, as demonstrated by increased BrdU+/HuC/D+ newborn neurons. The observed increase in microglial abundance and apoptotic cells suggests that NCBP2-AS2 influences both neurogenesis and cellular turnover. These results align with the critical role of energy metabolism in neural stem cell activity, particularly the balance between oxidative phosphorylation and glycolysis (Petrelli et al., 2023, Namba et al., 2021). The age-dependent effects on astroglial proliferation and consistent neurogenic increases highlight the regulatory role of NCBP2-AS2 in brain homeostasis. Notably, the zebrafish model provides a valuable platform to investigate these processes *in vivo*, facilitating the study of mitochondrial microproteins in neural contexts.

NCBP2-AS2 expression was significantly reduced in both human AD brains and zebrafish amyloid models. This conserved downregulation across species strengthens the link between NCBP2-AS2 and AD pathophysiology, highlighting its potential role in a shared mechanism underlying mitochondrial dysfunction and neurodegenerative progression. In amyloid-treated zebrafish, NCBP2-AS2 KO led to increased astroglial proliferation and neurogenesis, possibly representing a compensatory response to amyloid toxicity. These findings support the hypothesis that NCBP2-AS2 regulates neurogenic processes and that its downregulation may mitigate the effects of mitochondrial dysfunction in AD. Future research should investigate whether modulating NCBP2-AS2 expression in AD models can ameliorate neurodegenerative pathology.

### Strengths and limitations

This study presents significant strengths that advance the understanding of mitochondrial microproteins and their role in cellular physiology. Foremost, we identified NCBP2-AS2 as an evolutionarily conserved mitochondrial microprotein, underscoring its fundamental importance across species. Our findings bridge the gap between mitochondrial protein transport and energy metabolism, highlighting NCBP2-AS2 as a potential regulator of ATPase subunit translocation. We developed robust experimental tools that provide a strong foundation for further research. These include zebrafish and HEK293T knockout models, a stoichiometric co-immunoprecipitation platform, and metabolic assays, all of which enabled detailed functional analyses of NCBP2-AS2. Moreover, our use of electron microscopy and advanced interaction studies elucidated NCBP2-AS2’s localization and dynamic roles within the mitochondria. The integration of these tools not only strengthens our study but also offers resources for the scientific community to explore mitochondrial function in health and disease. However, limitations remain. We lack detailed structural information on NCBP2-AS2 and its interactions with mitochondrial complexes. Techniques such as crystallography or cryo-EM are needed to uncover its binding mechanisms with TIM chaperones and other partners. Furthermore, while we employed a single epitope tag for localization, the addition of complementary tags could enhance the detection of transient or weak interactions. Multi-omics approaches and longitudinal studies in human tissues are critical next steps to better understand NCBP2-AS2’s role in mitochondrial homeostasis and its implications in AD pathophysiology. Future research could explore the therapeutic potential of NCBP2-AS2 in addressing mitochondrial dysfunction and neurodegeneration, particularly through targeted interventions to modulate its activity.

## Materials and Methods

### Ethics statement

Zebrafish at various ages were used in this study (Landesdirektion Sachsen, Germany permit number: TVV32/2018). The fish were maintained under standard conditions as described (Geisler et al., 2016, Alestrom et al., 2019). All the procedures involved in animals were carried out in accordance with recommendations and official guidelines for laboratory animal handling and research and adhered to the white papers of European Zebrafish research community (Alestrom et al., 2019, Kohler et al., 2017). All efforts were made to avoid or minimize suffering.

### BAC recombination and generation of HeLa reporter lines

Bacterial artificial chromosome (BAC) recombination was performed by inserting the C-terminal LAP- cassette, developed by the MitoCheck Project, in frame into the NCBP2-AS2 gene by replacing its endogenous stop codon. Generation of plasmids, BAC modification, and transfections were performed by using the established protocols (Poser et al., 2008). The strategy includes three components: the LAP-cassette with Kanamycin-resistance gene to be integrated into the BAC with homology arms in its 5’ and 3’ end to human NSBC2-AS2 gene, pSC101 recombination plasmid with Tetracyline-resistance gene encodes the components of the Red/ET recombination system under the control of an arabinose- inducible promoter (GeneBridges), and the starting bacterial strains that contain the target BAC that contains chloramphenicol resistance (**Supplementary Data 1**). Bacterial cells containing the recombination plasmid pSC101 and the target BAC were grown overnight in the presence of chloramphenicol (28 ug/ml) and tetracycline (10 ug/ml) at the permissive temperature of 30 °C to allow for the propagation of the BAC and the pSC101 plasmid. The next morning these bacteria were induced with 0.2 % arabinose to activate the expression of the recombination proteins and incubated at 37 °C in the absence of tetracycline to stop the replication of the pSC101. Bacteria were electroporated with the purified LAP-cassette PCR product with the homology arms and afterwards the transformants were grown at 37 °C overnight in the presence of chloramphenicol and kanamycin (28 ug/ml for both) on agar plates to select for bacteria, where the integration of the LAP cassette into the BAC has occurred.

A colony PCR was used to identify colonies with the correct integration of the cassette. The primers were complementary to the genomic sequences upstream and downstream of the integration site, but outside of the homology regions used for Red/ET-mediated integration (**Supplementary Data 1**). The corresponding untagged BACs served as negative controls. PCR products were checked on an agarose gel and the positive colonies were then used to inoculate cultures at 37°C in medium containing chlorampenicol and kanamycin (or hygromycin and kanamycin). Cells were harvested by centrifugation and the tagged BACs were purified using a NucleoBond® PC20 Plasmid DNA Purification Kit according to the manufacturer’s instructions (Macherey-Nagel). After sequencing, the tagged BACs were introduced into HeLa cells by transfection as described (Poser et al., 2008). The LAP-cassette contains a geneticin/neomycin (G418) resistance gene for selection of mammalian cells. After the geneticin selection was over, the newly established BAC lines were sorted to increase the percentage of GFP- positive cells, and used for functional studies. The following product was used for selection of HeLa and MCF7 cells transfected with modified BACs: Geneticin 50 mg/ml (Gibco, 10131-019). For modifying the NCBP2-AS2 locus, the BAC MCB22042 (HS.139.A23) was provided by Dr. Susanne Hasse (TRansgeneOmics Facility, MPI-CBG, Dresden, Germany).

### Immunofluorescence on cells

Cells were seeded in Ibidi chambers in standard growth medium (+G418 for BAC-lines), with the desired cell numbers to achieve 80-90% confluence on the day of the analysis. Immunofluorescent detection was performed as described (Celikkaya et al., 2019, Papadimitriou et al., 2018). Images were acquired on the Delta Vision Elite Microscope (Applied Precision) by using the 60 X (Olympus, PlanApoN, 60X, 1.42, Oil) or 100 X (Olympus, UPlanSApo, 100X, 1.40, Oil) objectives, an immersion oil with refractive index of 1.518 (Applied Precision), and the InsightSSI Filter Set. The resulting images were deconvolved in softWoRx and adjustment of brightness and contrast were performed in FIJI. Antibodies used are listed in **Table S1**.

### Mitochondrial isolation and Western blotting

Mitochondrial isolation was performed by using the ab110171 kit (Abcam) following the manufacturer’s instructions. Protein concentration was measured using the Pierce 660 nm kit. Total protein extracts were mixed 1:1 with 2X Laemmli buffer, followed by boiling at 95°C for 5 min. The polyacrylamide gel electrophoresis (PAGE) was performed by using the XCell SureLock® Mini-Cell chamber and 10, 12 or 15 well 4-12% NuPAGE® Bis-Tris Mini Gels, the runnning buffer was 1X MOPS (Invitrogen). The nitrocellulose membrane was Amersham Protran 0.45 NC. Imaging of Western blot membranes was performed on the Odyssey Infrared Imaging System (LI-COR Biosciences), and adjustment of brightness and contrast were performed in the Image Studio Lite Analysis Software (LI-COR Biosciences).

### Electron microscopy

Cells were fixed fixed in 4% formaldehyde (prepared from para-formaldehyde prills) diluted in 100 mM Sörensen phosphate buffer (PB, pH 7.4) and processed for Tokuyasu cryo-sectioning and immunogold labeling as described^32,33^. In brief, the samples were washed in PB, infiltrated into 10% gelatin (1% for 30 min, 3% for 45 min, 7% for 1h, 10% for 2h) at 37°C, cooled down on ice, cut into small blocks (block size about 0.5 mm), incubated in 2.3 M sucrose in water for 24 hours at 4°C, mounted on pins (Leica # 16701950), and plunge frozen in liquid nitrogen. Ultrathin sections (70-100 nm) were prepared with a Leica UC6 ultramicrotome equipped with a FC6 cryo-chamber (Leica Microsystems, Wetzlar, Germany). The sections were picked up with methyl cellulose / sucrose (1 part 2% methyl cellulose (MC), Sigma M- 6385, 25 centipoises + 1 part 2.3 M sucrose) using a perfect loop and transferred to formvar-coated mesh grids. The sections were stained with antibodies against GFP (rabbit ant-GFP, Torrey Pines, # TP401, 1:100) and protein A 10 nm gold (UMC Utrecht, 1:50). For this, the grids were placed upside down on drops of PBS in a 37°C-incubator for 20 min to remove gelatin, sucrose, and methyl cellulose, then washed with 0.1% glycin / PBS (5x 1 min), blocked with 1% BSA/PBS (2x 5 min) and incubated with the primary antibodies for 1 hour. The sections were washed in PBS (4x 2 min) followed by incubation with protein A conjugated to 10 nm gold for 1 hour, washed again in PBS (3x 5 s, 4x 2 min) and post-fixed in 1% glutaraldehyde (GA, 5 min). After several washes in water the sections were contrasted with 1% aqueous uranyl acetate. Contrasted sections were analyzed on a Jeol JEM1400 Plus (JEOL, Freising, Germany) running at 80 kV acceleration voltage and equipped with a digital camera (Ruby, Jeol).

### Generation of CRISPR knockout for NCBP2-AS2 in HEK293T cell lines

The following gRNA sequences were designed by using CRISPR Scan algorithm (koAS2gR1F: CACCGCTTGGAGAAGATGGTGCTGCGG; koAS2gR1R: AAACCCGCAGCACCATCTTCTCCAAGC).

The gRNA sequences were cloned into the PX458 vector, which contains the coding sequence of the Cas9 enzyme and a GFP reporter by using a previously published protocol (Ran et al., 2013). Plasmid DNA was purified with the Thermo Scientific GeneJET Plasmid Maxiprep Kit (K0492) and sequenced (Eurofins). HEK293T cells were transfected with the PX458 plasmid, containing the cloned gRNA, or with the starting “empty” PX458 plasmid. Single GFP-positive cells were sorted into wells of a 96-well plate. Surviving clones were grown and DNA from pellets of 1 million cells were prepared for DNA analysis. The target region of deletion was PCR amplified using the following primers: Reverse: CCCTCCACATTGATGCCACT, forward: GAGGCGAGCATCTCATTGGA. T7 Endonuclease I analyses were performed on the denatured and re-annealed duplex PCR products to validate the modifications according to the manufacturer’s instructions (NEB). In the case of small insertions or deletions, a stretch of mismatched bases is formed after annealing, which is a substrate for the T7 endonuclease I enzyme that cleaves at such sites. The presence of the expected restriction pattern on the agarose gel shows that the desired genomic modification occurred in the transfected cells. Changes of expression was determined by qRT-PCR (**Data S2**).

### Expression of zebrafish NCBP2-AS2 in HEK cells

Zebrafish and human NCBP2-AS2 were cloned into pIRES-EGFP plasmid (6029-1, Clontech) for expression in human cells and localization analyses. The transfection was carried out according to the Lipofectamine® 2000 DNA Transfection Reagent Protocol. HEK293T cell line was seeded at the density of 50,000 cells per well in a Nunc™ Cell-Culture Treated 24-well plate on high precision cover slips (170 μm ± 5 μm) coated with 0,1% gelatin solution. The cells were cultured in Dulbecco’s Modified Eagle Medium (DMEM) supplemented with 10% Fetal Bovine Serum (FBS) and 1% Penicillin-Streptomycin (P/S). One day after seeding, the cells were transfected with 500 ng of pIRES2-EGFP (Clontech), hAS2- Flag-Strep-pIRES2-EGFP and zfAS2-Flag8 Strep-pIRES2-EGFP (Supplementary Data 1) and 3 μl Lipofectamine. A medium change was performed one day post transfection. Two days post transfection the cells were fixed with 4% prewarmed Paraformaldehyde (PFA) for 10 minutes at room temperature. Fixed cells were permeabilized with 100 mM digitonin (1:1000, final 100 uM) in PBS for five minutes and then washed once in PBS. The blocking was performed with 2% Bovine Serum Albumin (BSA) in PBS for one hour. The cells were incubated over night at 4 ℒC with primary antibody solution with anti-TOM20 and anti-FLAG or PAM16 and anti-FLAG-M2 in 2% BSA in PBS. Further, the cells were incubated for 1 hour at room temperature with secondary antibody solution with Goat anti-mouse and Goat anti-rabbit or Goat anti-mouse and Goat anti-rabbit. Nuclear staining was performed with 100 mg/ml DAPI. This was followed by a post-fixation step with 4% PFA. The cover slips were then mounted on glass slides with ProLong™ Glass Antifade Mountant P36982. The images were acquired with Zeiss LSM 780 NLO system with a 100x objective.

### Co-immunoprecipitation

For proteomics, hAS2 and control plasmids have been transfected into knock-out HEK cells in varying equimolar ratios (Figure 2), followed by protein affinity purification (beads) and western blot. By using this strategy, one can make sure all of the target proteins (hsAS2) are actually tagged (in our case by Flag), on a knockout background, because in principle the untagged endogenous version of hsAS2 cannot be pulled down with the anti-Flag beads but will compete with the exogenous tagged version for the interaction partners. The mixture of control and experimental protein lysates into one tube for co-IP (volume-wise) allows one to distinguish true interactors from contaminants (the true interactors should be high in the co-IP mixture with highest percentage of the experimental lysate, and the interactors should be diluted when more and more control lysate is added to the co-IP mixtures, while the contaminants will be more consistent across all mixtures and altogether have a different behavior).

Four million HEK293T cells where plated on 10 cm culture coated dishes (4 plates in total). PEI transfection was performed the following day by adding the PEI complex to DNA complex (polyethylenimine (PEI) stock solution (1 mg/ml PEI in PBS)). Following the overnight incubation, medium was changed to DMEM with 10% FBS and 1%P/S. Protein purification was performed at two days post transfection. Cells were incubated in HBSNO buffer with protease and phosphatase inhibitors (1 tablet each in 10 ml), as well as RNaseA and DNaseI (10 mM HEPES-KOH, pH 7.5, 150 mM NaCl, 0.5% NP-40, 0.5% OGP, 5% Glycerol, supplemented freshly before use with 1 ug/ml RNAseA, 1 ug/ml DNAse I (1:100 from stocks), complete protease inhibitor, PhosStop phosphatase inhibitor). Experimental and control cells were washed twice with room temperature PBS in their dishes (2 dishes per group – one group transfected with the control plasmid, the other group transfected with the hsAS2 overexpression plasmid, **Data S1**). Cells were lysed and lysates were subjected to BCA assay. ANTI- FLAG M2 Affinity Gel (A2220) was prepared and agarose beads (180 ul bead suspension for 4 samples) with 1800 ul (10 volumes) of supplemented HBSNO were pre-washed. Bead suspension was added to the lysates and incubated for 30-45 min min at 4 C. Beads were collected by short centrifugation at 1000 g for 1 minutes at 4 C. Beads were washed gently in column once with supplemented HBSNO, and twice with HBS/PhosStop (200 ul each, 5 min, cold room; 10 mM HEPES-KOH pH 7.5, 150 mM NaCl, supplemented freshly before use with PhosStop phosphatase inhibitor). Interaction complexes were eluted with 40 ul 100 mM Glycine. Pulldown samples were transferred to -20 C until further analysis by MS. Western Blot was performed following the instruction of NuPAGE® Technical guide. The samples were run on a NuPAGE® Novex® Bis-Tris Gel with 1xMES SDS Running Buffer at 200 V for 50 minutes and transferred on a PVDF membrane at 30V for 1h. Transfer buffer contains 20% Methanol. The membrane was blocked for 1 hour at room temperature in 5% Milk solution and incubated over night at 4 ℒC with primary antibody solution (anti-FLAG-M2, mouse and anti-GFP, rabbit 1:1000). Incubation with secondary antibody was performed for 1 hour at room temperature. The images were captured with Odyssey® CLx Imaging System. For reagent details see **Table S1**.

### Proteomics and LC-MS/MS

The samples were digested On-Beads in solution with subsequently twice with 200ng trypsin (Promega) for 18 hrs and finally with 50 ng rLys-C (Promega) for 24 hrs. The digestion solutions were desalted with C18 ultramicro-columns (Nest Group, USA), eluted with 50% acetonitrile, 0.1 % formic acid and the eluates were dried under vacuum and stored until analysis at -20 °C. For LC-MS/MS analyses the peptides were recovered in 5 µl 30 % formic acid supplemented with 25 fmol/µl Peptide Retention Time Standard (ThermoScientific – Pierce), diluted with 20 µl of water and 5 µl were injected. NanoLC-MS/MS analyses were performed with a Q-Exactive HF mass spectrometer (ThermoScientific) hyphenated to nanoflow LC system (Dionex3000 RSLC, ThermoScientific). Peptides were separated in linear gradient of 0.1 % aqueous formic acid (eluent A) and 0.1 % formic acid in 60% acetonitrile (eluent B) and the mass spectrometer was operated in data-dependent acquisition mode (DDA, TopN 10). Raw files were loaded into the Progenesis QIP V4.2 software (Nonlinear Dynamics, Newcastle u.T., UK) for peak picking and quantitative analysis by MI3/HI3 (Silva et al., 2006). Peptide and protein identification was performed with Mascot V2.6 (Matrixscience, UK). The proteomics data have been deposited at the PRIDE database (Perez-Riverol et al., 2019), member of the ProteomeXchange (Deutsch et al., 2020) consortium for proteomic data (EBI, UK) under the accession PXD041213, **Data S3**). Detailed parameters are listed in **Table S2**. For validation of proteomics hits, we investigated the levels of SLC25A6, C190rf52, TIMM9, ATP5F1C and SLC25A6 by using MTCO1 as normalization control in HEK293T cells with or without NCBP2-AS2 knock-out with the western blot procedure described above.

### Metabolic analyses of cell lines

To determine the carbon metabolic rates of HEK293T cells with or without NCBP2-AS2 knockout, we used mammalian cell metabolic assay kit (Catalog numbers 13101-13104, 72301, 74352, Biolog). Cultured cells were count-adjusted as instructed by the manufacturer on provided plates with select substrates. Redox dye treatment and kinetic measurements were performed every 30 mins for 5 hours. Calibration, normalization, and plotting was performed as per manufacturer’s instructions. Raw data in **Data S4**.

### RNA extraction, cDNA synthesis, RT-PCR, single cell sequencing

Zebrafish telencephalon was dissected, RNA was isolated, and cDNA was generated as described (Siddiqui et al., 2021, Chang et al., 2021, Bhattarai et al., 2020, Bhattarai et al., 2016). For primers see **Table S1**. Single cell sequencing data was used from previous published datasets and tSNE plots were generated as described (Fore et al., 2020, Cosacak et al., 2020b, Cosacak et al., 2019).

### Generation of NCBP2-AS2 knockout with CRISPR in zebrafish

The following gRNA sequences were used to generate DNA templates for in vitro transcription of guide RNAs for zebrafish NCBP2-AS2 gene (ENSDARG00000104726): zf_AS2_gRNA5:taatacgactcactataGGGCTGTTAATGAGAAAGTGgttttagagctagaa, zf_AS2_gRNA6_Cter_79: taatacgactcactata GGGAGGTCCGAGAGGGAGTGgttttagagctagaa and gRNA_universal_tail_R:AAAAGCACCGACTCGGTGCCACTTT TTCAAGTTGATAACGGACTAGCCTTATTTTAACTTGCTATTTCTAGCTCTAAAAC. Universal tail primer was separately used with reverse and forward primer to generate DNA templates, with flanking gRNA sites. PCR product was purified (PCR purification kit, Qiagen) and concentration was measured with NanoDrop. For in vitro transcription MEGAshortscript kit (Ambion) was used as per manufacturer’s instructions. Purification was performed as described (Kizil et al., 2009). Cas9 RNA was prepared from the vector deposited to AddGene by Wenbiao Chen lab (plasmid 46757, pT3TS-nCas9n; (Jao et al., 2013)) using mMESSAGE mMACHINE kit (Ambion). 600 pg Cas9 mRNA, 100 pg gRNA1 and gRNA2 were injected into 1-cell stage embryos using Pneumatic Picopump PV820 (World Precision Instruments) with TW100F-3 glass capillaries with filament (OD/ID 1/0.75 mm). Needles were prepared with Sutter Instrument (Vel 50, Time 80, Pull 250, Heat 253). Fin clipping and PCR, followed by agarose gel electrophoresis was performed as described (Nüsslein-Volhard and Dahm, 2002) to identify the founder zebrafish for their genotype status: wild-type, heterozygous deletion and homozygous knock-out (for primers see **Table S1**). Experiments were performed with F3 generation from sequence-verified knockout animals.

### Phenotypic analyses of the effects of NCBP2-AS2 deletion in adult zebrafish telencephalon

Dissection of zebrafish telencephalon (wild type or homozygous deletion of NCBP2-AS2), fixation, tissue preparation, embedding, histological sectioning, immunostaining, imaging, and quantifications were performed as established and described before (Lee et al., 2022, Siddiqui et al., 2021, Bhattarai et al., 2020, Bhattarai et al., 2016). For the quantification of newborn neurons, the fish were treated with 10 uM Bromodeoxyuridine (BrdU) for 7 hrs each in two consecutive days and were supported with the air- interchange system. After 14 days, fish were euthanized. For the analysis of stem cell proliferation, the imaging was done automatically by Zeiss ApoTome Axioimager.Z1, and the PCNA/S100 double-positive cells were quantified by manual counting via ZEN 2012 Blue. For the analysis of inflammatory response, the quantification was carried out by live manual counting through Zeiss ApoTome.2. The representative images were taken through Zeiss ApoTome.2 and processed via ZEN 2012 Blue. For the analysis of apoptosis, the imaging was carried out by Zeiss ApoTome.2, and the quantification of TUNEL positive cells was done by manual counting via ZEN 2012 Blue. Manual counting was performed blindly. Statistical analyses were performed with GraphPad Prism 8.0 analysis software. Unpaired nonparametric t test with Kolmogorov-Smirnov test for pairwise comparisons. Conventional * nomenclature used. (*: p<0.05, **: p<0.01 or ***: p<0.005).

### Transcriptomics data analysis

Single-cell/nucleus sequencing datasets were obtained from previously published studies (Tayran et al., 2024, Is et al., 2024). For both datasets, raw count matrices and associated metadata were downloaded from public repositories. Data preprocessing, including quality control, normalization, and clustering, was performed using the Seurat package (version 4.0) in R. Cells with fewer than 200 detected genes or greater than 10% mitochondrial gene content were excluded to remove low-quality cells. Normalized expression data were integrated and scaled prior to principal component analysis (PCA) for dimensionality reduction. Uniform Manifold Approximation and Projection (UMAP) was applied to visualize clustering and assign cell types based on marker gene expression. Annotation of cell types was confirmed by matching cluster-specific marker genes with established cell type markers for both zebrafish and human datasets. For differential gene expression analysis, the Wilcoxon rank-sum test was employed to identify genes with significant differences between experimental conditions. Expression levels were calculated as the average expression across all clusters corresponding to each specific cell type.

### Author contributions

Conceptualization: SP, CK; Methodology: SP, PB, EY, AS, TK, FB, MG, CK; Primary data acquisition: SP, PB, DL, JHK, GE, BC, JM, MIC, MG, TK; Data analyses: SP, DL, JHK, EY, MIC, MG; Funding: FB, CK; Resources and reagents: AS, MG, TK, FB, CK. Writing, editing and revisions: all authors.

## Acknowledgements

We would like to thank Kristin Eismann (Core Facility Mass Spectrometry & Proteomics, CMCB, TU Dresden), and Kerstin Brandt and Heike Hollak (DZNE Dresden, Helmholtz Association) for technical assistance; Michael Redd for L-Plastin antibody; Wenbiao Chen lab for Cas9 plasmid; and CMCB/CRTD TU Dresden core facilities (imaging and electron microscopy). This work was supported by German Center for Neurodegenerative Diseases core funding (C.K.), Helmholtz Young Investigator Award (C.K.), and Deutsche Forschungsgemeinschaft individual research grant (C.K.), Columbia University Schaefer Research Scholars Program Award (C.K), National Institute on Aging R01 AG067501 (C.K., PD/PI: Richard Mayeux) and National Institute on Aging RF1 AG066107 (C.K., PD/PI: Richard Mayeux), Erasmus mobility program funding (G.E., B.C.) and Graduate Academy of TU Dresden fellowship (S.P.). The CF Mass Spectrometry & Proteomics was supported by grants from the German Federal Ministry of Education and Research (BMBF) program ’Unternehmen Region’ (grants # 03Z2ES1, # 03Z22EB1), the German Research Foundation (INST 269/731-1 FUGG) and the European Regional Development Fund (ERDF/EFRE) (Contract # 100232736).

## Supplementary Materials

**Supplementary Data 1:** GenBank files for the plasmids used for this study.

**Supplementary Data 2:** qRT PCR measurement raw data for the NCBP2-AS2 expression in knockout HEK293T cell clones.

**Supplementary Data 3:** Co-immunoprecipitation and LC-MS results.

**Supplementary Data 4:** Western blots and quantification for selected candidates.

**Supplementary Data 5:** Metabolic assay raw reads and activity measurements.

**Supplementary Data 6:** Quantification data for zebrafish.

**Supplementary Table 1:** List of materials, reagents, and equipment.

**Supplementary Table 2:** Mass spectroscopy and liquid chromatography setttings and specifications.

## References

1. Alestrom, P., D’angelo, L., Midtlyng, P. J., Schorderet, D. F., SCHULTE-Merker, S., Sohm, F. & Warner, S. 2019. Zebrafish: Housing and husbandry recommendations. Lab Anim, 23677219869037.

2. Anderson, D. M., Anderson, K. M., Chang, C. L., Makarewich, C. A., Nelson, B. R., Mcanally, J. R., Kasaragod, P., Shelton, J. M., Liou, J., BASSEL-Duby, R. & Olson, E. N. 2015. A micropeptide encoded by a putative long noncoding RNA regulates muscle performance. Cell, 160, 595–606.

3. Askenazi, M., Kavanagh, T., Pires, G., Ueberheide, B., Wisniewski, T. & Drummond, E. 2023. Compilation of reported protein changes in the brain in Alzheimer’s disease. Nat Commun, 14, 4466.

4. Bai, B., Vanderwall, D., Li, Y., Wang, X., Poudel, S., Wang, H., Dey, K. K., Chen, P. C., Yang, K. & Peng, J. 2021. Proteomic landscape of Alzheimer’s Disease: novel insights into pathogenesis and biomarker discovery. Mol Neurodegener, 16, 55.

5. Benezra, R., Davis, R. L., Lassar, A., Tapscott, S., Thayer, M., Lockshon, D. & Weintraub, H. 1990. Id: a negative regulator of helix-loop-helix DNA binding proteins. Control of terminal myogenic differentiation. Ann N Y Acad Sci, 599, 1-11.

6. Beverly, K. N., Sawaya, M. R., Schmid, E. & Koehler, C. M. 2008. The Tim8-Tim13 complex has multiple substrate binding sites and binds cooperatively to Tim23. J Mol Biol, 382, 1144–56.

7. Bhattarai, P., Cosacak, M. I., Mashkaryan, V., Demir, S., Popova, S., Govindarajan, N., Brandt, K., Zhang, Y., Chang, W., Ampatzis, K. & Kizil, C. 2020. Neuron-glia interaction through Serotonin-BDNF-NGFR axis enables regenerative neurogenesis in Alzheimer’s model of adult zebrafish brain. . PLoS Biol, 18, e3000585.

8. Bhattarai, P., Gunasekaran, T. I., Belloy, M. E., Reyes-Dumeyer, D., Julich, D., Tayran, H., Yilmaz, E., Flaherty, D., Turgutalp, B., Sukumar, G., Alba, C., Mcgrath, E. M., Hupalo, D. N., Bacikova, D., LE Guen, Y., Lantigua, R., Medrano, M., Rivera, D., Recio, P., Nuriel, T., Ertekin-Taner, N., Teich, A. F., Dickson, D. W., Holley, S., Greicius, M., Dalgard, C. L., Zody, M., Mayeux, R., Kizil, C. & Vardarajan, B. N. 2024. Rare genetic variation in fibronectin 1 (FN1) protects against APOEepsilon4 in Alzheimer’s disease. Acta Neuropathol, 147, 70.

9. Bhattarai, P., Thomas, A. K., Cosacak, M. I., Papadimitriou, C., Mashkaryan, V., Zhang, Y. & Kizil, C. 2017a. Modeling Amyloid-β42 Toxicity and Neurodegeneration in Adult Zebrafish Brain. Journal of Visualized Experiments, 128.

10. Bhattarai, P., Thomas, A. K., Papadimitriou, C., Cosacak, M. I., Mashkaryan, V., Froc, C., Kurth, T., Dahl, A., Zhang, Y. & Kizil, C. 2016. IL4/STAT6 signaling activates neural stem cell proliferation and neurogenesis upon Amyloid-β42 aggregation in adult zebrafish brain. Cell Reports, 17, 941–948.

11. Bhattarai, P., Thomas, A. K., Zhang, Y. & Kizil, C. 2017b. The effects of aging on Amyloid-β42- induced neurodegeneration and regeneration in adult zebrafish brain. Neurogenesis, 4, e1322666.

12. Bhattarai, P., Turgutalp, B. & Kizil, C. 2022. Zebrafish as an Experimental and Preclinical Model for Alzheimer’s Disease. ACS Chem Neurosci, 13, 2939–2941.

13. Celikkaya, H., Cosacak, M. I., Papadimitriou, C., Popova, S., Bhattarai, P., Biswas, S. N., Siddiqui, T., Wistorf, S., Nevado-Alcalde, I., Naumann, L., Mashkaryan, V., Brandt, K., Freudenberg, U., Werner, C. & Kizil, C. 2019. GATA3 Promotes the Neural Progenitor State but Not Neurogenesis in 3D Traumatic Injury Model of Primary Human Cortical Astrocytes. Front Cell Neurosci, 13, 23.

14. Chang, W., Pedroni, A., Bertuzzi, M., Kizil, C., Simon, A. & Ampatzis, K. 2021. Locomotion dependent neuron-glia interactions control neurogenesis and regeneration in the adult zebrafish spinal cord. Nat Commun, 12, 4857.

15. Cosacak, M. I., Bhattarai, P., DE Jager, P. L., Menon, V., Tosto, G. & Kizil, C. 2022. Single Cell/Nucleus Transcriptomics Comparison in Zebrafish and Humans Reveals Common and Distinct Molecular Responses to Alzheimer’s Disease. Cells, 11, 1807.

16. Cosacak, M. I., Bhattarai, P. & Kizil, C. 2020a. Alzheimer’s disease, neural stem cells and neurogenesis: cellular phase at single-cell level. Neural Reg Res, 15, 824–827.

17. Cosacak, M. I., Bhattarai, P. & Kizil, C. 2020b. Protocol for Dissection and Dissociation of Zebrafish Telencephalon for Single-Cell Sequencing. STAR Protoc, 1, 100042.

18. Cosacak, M. I., Bhattarai, P., Reinhardt, S., Petzold, A., Dahl, A., Zhang, Y. & Kizil, C. 2019. Single-Cell Transcriptomics Analyses of Neural Stem Cell Heterogeneity and Contextual Plasticity in a Zebrafish Brain Model of Amyloid Toxicity. Cell Rep, 27, 1307–1318 e3.

19. Deutsch, E. W., Bandeira, N., Sharma, V., Perez-Riverol, Y., Carver, J. J., Kundu, D. J., Garcia-Seisdedos, D., Jarnuczak, A. F., Hewapathirana, S., Pullman, B. S., Wertz, J., Sun, Z., Kawano, S., Okuda, S., Watanabe, Y., Hermjakob, H., Maclean, B., Maccoss, M. J., Zhu, Y., Ishihama, Y. & Vizcaino, J. A. 2020. The ProteomeXchange consortium in 2020: enabling ’big data’ approaches in proteomics. Nucleic Acids Res, 48, D1145–D1152.

20. Felsky, D., Santa-Maria, I., Cosacak, M. I., French, L., Schneider, J. A., Bennett, D. A., DE Jager, P. L., Kizil, C. & Tosto, G. 2023. The Caribbean-Hispanic Alzheimer’s disease brain transcriptome reveals ancestry-specific disease mechanisms. Neurobiol Dis, 176, 105938.

21. Fore, S., Acuña-Hinrichsen, F., Mutlu, K. A., Bartoszek, E. M., Serneels, B., Faturos, N. G., Chau, K. T. P., Cosacak, M. I., Verdugo, C. D., Palumbo, F., Ringers, C., JURISCH-Yaksi, N., Kizil, C. & Yaksi, E. 2020. Functional properties of habenular neurons are determined by developmental stage and sequential neurogenesis. Science Advances, 6, eaaz3173.

22. Geisler, R., Borel, N., Ferg, M., Maier, J. V. & Strahle, U. 2016. Maintenance of Zebrafish Lines at the European Zebrafish Resource Center. Zebrafish, 13 Suppl 1, S19–23.

23. Gray, L. R., Tompkins, S. C. & Taylor, E. B. 2014. Regulation of pyruvate metabolism and human disease. Cell Mol Life Sci, 71, 2577–604.

24. Grevel, A. & Becker, T. 2020. Porins as helpers in mitochondrial protein translocation. Biol Chem, 401, 699–708.

25. Haage, V., Tuddenham, J. F., Comandante-Lou, N., Bautista, A., Monzel, A., Chiu, R., Fujita, M., Garcia, F. G., Bhattarai, P., Patel, R., Buonfiglioli, A., Idiarte, J., Herman, M., Rinderspacher, A., Mela, A., Zhao, W., Argenziano, M. G., Furnari, J. L., Banu, M. A., Landry, D. W., Bruce, J. N., Canoll, P., Zhang, Y., Nuriel, T., Kizil, C., Sproul, A. A., DE Witte, L. D., Sims, P. A., Menon, V., Picard, M. & De Jager, P. L. 2024. A pharmacological toolkit for human microglia identifies Topoisomerase I inhibitors as immunomodulators for Alzheimer’s disease. bioRxiv.

26. Hassel, K. R., Brito-Estrada, O. & Makarewich, C. A. 2023. Microproteins: Overlooked regulators of physiology and disease. iScience, 26, 106781.

27. Ikeda, K., Shiba, S., Horie-Inoue, K., Shimokata, K. & Inoue, S. 2013. A stabilizing factor for mitochondrial respiratory supercomplex assembly regulates energy metabolism in muscle. Nat Commun, 4, 2147.

28. Is, O., Wang, X., Reddy, J., Min, Y., Yilmaz, E., Bhattarai, P., Patel, T., Bergman, J., Quicksall, Z., Heckman, M., Tutor-New, F., Demirdöğen, B. C., White, L., Koga, S., Krause, V., Inoue, Y., Kanekiyo, T., Cosacak, M. I., Nelson, N., Lee, A., Vardarajan, B., Mayeux, R., Kouri, N., Deniz, K., Carnwath, T., Oatman, S., Lewis-Tuffin, L., Nguyen, T., Carrasquillo, M., Graff-Radford, J., Petersen, R. C., Jack Jr, C., Kentarci, K., Murray, M., Nho, K., Saykin, A., Dickson, D., Kizil, C., Allen, M. & Ertekin-Taner, N. 2024. Gliovascular transcriptional perturbations in Alzheimer’s disease reveal molecular mechanisms of blood brain barrier dysfunction. Nat Commun, 15, 4758.

29. Jao, L. E., Wente, S. R. & Chen, W. 2013. Efficient multiplex biallelic zebrafish genome editing using a CRISPR nuclease system. Proc Natl Acad Sci U S A, 110, 13904–9.

30. Kalia, V., Niedzwiecki, M. M., Bradner, J. M., Lau, F. K., Anderson, F. L., Bucher, M. L., Manz, K. E., Schlotter, A. P., Fuentes, Z. C., Pennell, K. D., Picard, M., Walker, D. I., Hu, W. T., Jones, D. P. & Miller, G. W. 2022. Cross-species metabolomic analysis of tau- and DDT-related toxicity. PNAS Nexus, 1, pgac050.

31. Kang, Y., Baker, M. J., Liem, M., Louber, J., Mckenzie, M., Atukorala, I., Ang, C. S., Keerthikumar, S., Mathivanan, S. & Stojanovski, D. 2016. Tim29 is a novel subunit of the human TIM22 translocase and is involved in complex assembly and stability. Elife, 5.

32. Kawatani, K., Holm, M. L., Starling, S. C., Martens, Y. A., Zhao, J., Lu, W., Ren, Y., Li, Z., Jiang, P., Jiang, Y., Baker, S. K., Wang, N., Roy, B., Parsons, T. M., Perkerson, R. B., 3rd, Bao, H., Han, X., Bu, G. & Kanekiyo, T. 2023. ABCA7 deficiency causes neuronal dysregulation by altering mitochondrial lipid metabolism. Mol Psychiatry.

33. Kim, J. & Cheong, J. H. 2020. Role of Mitochondria-Cytoskeleton Interactions in the Regulation of Mitochondrial Structure and Function in Cancer Stem Cells. Cells, 9.

34. Kim, Y. A., Mellen, M., Kizil, C. & Santa-Maria, I. 2023a. Mechanisms linking cerebrovascular dysfunction and tauopathy: Adding a layer of epiregulatory complexity. Br J Pharmacol.

35. Kim, Y. A., Siddiqui, T., Blaze, J., Cosacak, M. I., Winters, T., Kumar, A., Tein, E., Sproul, A. A., Teich, A. F., Bartolini, F., Akbarian, S., Kizil, C., Hargus, G. & SANTA-Maria, I. 2023b. RNA methyltransferase NSun2 deficiency promotes neurodegeneration through epitranscriptomic regulation of tau phosphorylation. Acta Neuropathol, 145, 29–48.

36. Kizil, C. 2018. Mechanisms of Pathology-Induced Neural Stem Cell Plasticity and Neural Regeneration in Adult Zebrafish Brain. Current Pathobiology Reports, 6, 71–77.

37. Kizil, C. & Bhattarai, P. 2018. Is Alzheimer’s Also a Stem Cell Disease? - The Zebrafish Perspective. Front Cell Dev Biol, 6, 159.

38. Kizil, C., Otto, G. W., Geisler, R., Nusslein-Volhard, C. & Antos, C. L. 2009. Simplet controls cell proliferation and gene transcription during zebrafish caudal fin regeneration. Dev Biol, 325, 329–40.

39. Kizil, C., Sariya, S., Kim, Y. A., Rajabli, F., Martin, E., Reyes-Dumeyer, D., Vardarajan, B., Maldonado, A., Haines, J. L., Mayeux, R., JIMENEZ-Velazquez, I. Z., Santa-Maria, I. & Tosto, G. 2022. Admixture Mapping of Alzheimer’s disease in Caribbean Hispanics identifies a new locus on 22q13.1. Mol Psychiatry.

40. Kohler, A., Collymore, C., Finger-Baier, K., Geisler, R., Kaufmann, L., Pounder, K. C., SCHULTE- Merker, S., Valentim, A., Varga, Z. M., Weiss, J. & Strahle, U. 2017. Report of Workshop on Euthanasia for Zebrafish-A Matter of Welfare and Science. Zebrafish, 14, 547–551.

41. Kugeratski, F. G., Atkinson, S. J., Neilson, L. J., Lilla, S., Knight, J. R. P., Serneels, J., Juin, A., Ismail, S., Bryant, D. M., Markert, E. K., Machesky, L. M., Mazzone, M., Sansom, O. J. & Zanivan, S. 2019. Hypoxic cancer-associated fibroblasts increase NCBP2-AS2/HIAR to promote endothelial sprouting through enhanced VEGF signaling. Sci Signal, 12.

42. Lee, A. J., Raghavan, N. S., Bhattarai, P., Siddiqui, T., Sariya, S., Reyes-Dumeyer, D., Flowers, X. E., Cardoso, S. A. L., DE Jager, P. L., Bennett, D. A., Schneider, J. A., Menon, V., Wang, Y., Lantigua, R., Medrano, M., Rivera, D., Jimenez-Velazquez, I. Z., Kukull, W. A., Brickman, A. M., Manly, J. J., Tosto, G., Kizil, C., Vardarajan, B. & Mayeux, R. 2022. FMNL2 regulates gliovascular interactions and is associated with vascular risk factors and cerebrovascular pathology in Alzheimer’s disease. Acta Neuropathol.

43. Lin, Y. F., Xiao, M. H., Chen, H. X., Meng, Y., Zhao, N., Yang, L., Tang, H., Wang, J. L., Liu, X., Zhu, Y. & Zhuang, S. M. 2019. A novel mitochondrial micropeptide MPM enhances mitochondrial respiratory activity and promotes myogenic differentiation. Cell Death Dis, 10, 528.

44. Liu, L., Peritore, C., Ginsberg, J., Kayhan, M. & Donmez, G. 2015. SIRT3 attenuates MPTP-induced nigrostriatal degeneration via enhancing mitochondrial antioxidant capacity. Neurochem Res, 40, 600–8.

45. Makarewich, C. A., Baskin, K. K., Munir, A. Z., Bezprozvannaya, S., Sharma, G., Khemtong, C., Shah, A. M., Mcanally, J. R., Malloy, C. R., Szweda, L. I., BASSEL-Duby, R. & Olson, E. N. 2018a. MOXI Is a Mitochondrial Micropeptide That Enhances Fatty Acid beta-Oxidation. Cell Rep, 23, 3701–3709.

46. Makarewich, C. A., Munir, A. Z., Schiattarella, G. G., Bezprozvannaya, S., Raguimova, O. N., Cho, E. E., Vidal, A. H., Robia, S. L., BASSEL-Duby, R. & Olson, E. N. 2018b. The DWORF micropeptide enhances contractility and prevents heart failure in a mouse model of dilated cardiomyopathy. Elife, 7.

47. March-Diaz, R., Lara-Urena, N., Romero-Molina, C., Heras-Garvin, A., Ortega-De San Luis, C., Alvarez-Vergara, M., Sanchez-Garcia, M., Sanchez-Mejias, E., Davila, J., Rosales-Nieves, A., Forja, C., Navarro, V., Gomez-Arboledas, A., Sanchez-Mico, M., Viehweger, A., Gerpe, A., Hodson, E., Vizuete, M., Bishop, T., Serrano-Pozo, A., Lopez-Barneo, J., Berra, E., Gutierrez, A., Vitorica, J. & Pascual, A. 2021. Hypoxia compromises the mitochondrial metabolism of Alzheimer’s disease microglia via HIF1. Nature Aging, 1, 385–399.

48. Miller, B., Kim, S. J., Mehta, H. H., Cao, K., Kumagai, H., Thumaty, N., Leelaprachakul, N., Braniff, R. G., Jiao, H., Vaughan, J., Diedrich, J., Saghatelian, A., Arpawong, T. E., Crimmins, E. M., Ertekin-Taner, N., Tubi, M. A., Hare, E. T., Braskie, M. N., Decarie-Spain, L., Kanoski, S. E., Grodstein, F., Bennett, D. A., Zhao, L., Toga, A. W., Wan, J., Yen, K., Cohen, P. & ALZHEIMER’S DISEASE Neuroimaging, I. 2022. Mitochondrial DNA variation in Alzheimer’s disease reveals a unique microprotein called SHMOOSE. Mol Psychiatry.

49. Mor, D. E., Sohrabi, S., Kaletsky, R., Keyes, W., Tartici, A., Kalia, V., Miller, G. W. & Murphy, C. T. 2020. Metformin rescues Parkinson’s disease phenotypes caused by hyperactive mitochondria. Proc Natl Acad Sci U S A, 117, 26438–26447.

50. Namba, T., Nardelli, J., Gressens, P. & Huttner, W. B. 2021. Metabolic Regulation of Neocortical Expansion in Development and Evolution. Neuron, 109, 408–419.

51. Nüsslein-Volhard, C. & Dahm, R. 2002. Zebrafish: A Practical Approach, Oxford University Press.

52. Papadimitriou, C., Celikkaya, H., Cosacak, M. I., Mashkaryan, V., Bray, L., Bhattarai, P., Brandt, K., Hollak, H., Chen, X., He, S., Antos, C. L., Lin, W., Thomas, A. K., Dahl, A., Kurth, T., Friedrichs, J., Zhang, Y., Freudenberg, U., Werner, C. & Kizil, C. 2018. 3D Culture Method for Alzheimer’s Disease Modeling Reveals Interleukin-4 Rescues Abeta42-Induced Loss of Human Neural Stem Cell Plasticity. Dev Cell, 46, 85-101 e8.

53. Paschen, S. A., Rothbauer, U., Kaldi, K., Bauer, M. F., Neupert, W. & Brunner, M. 2000. The role of the TIM8-13 complex in the import of Tim23 into mitochondria. EMBO J, 19, 6392–400.

54. Perez-Riverol, Y., Csordas, A., Bai, J., Bernal-Llinares, M., Hewapathirana, S., Kundu, D. J., Inuganti, A., Griss, J., Mayer, G., Eisenacher, M., Perez, E., Uszkoreit, J., Pfeuffer, J., Sachsenberg, T., Yilmaz, S., Tiwary, S., Cox, J., Audain, E., Walzer, M., Jarnuczak, A. F., Ternent, T., Brazma, A. & Vizcaino, J. A. 2019. The PRIDE database and related tools and resources in 2019: improving support for quantification data. Nucleic Acids Res, 47, D442–D450.

55. Petrelli, F., Scandella, V., Montessuit, S., Zamboni, N., Martinou, J. C. & Knobloch, M. 2023. Mitochondrial pyruvate metabolism regulates the activation of quiescent adult neural stem cells. Sci Adv, 9, eadd5220.

56. Poser, I., Sarov, M., Hutchins, J. R., Heriche, J. K., Toyoda, Y., Pozniakovsky, A., Weigl, D., Nitzsche, A., Hegemann, B., Bird, A. W., Pelletier, L., Kittler, R., Hua, S., Naumann, R., Augsburg, M., Sykora, M. M., Hofemeister, H., Zhang, Y., Nasmyth, K., White, K. P., Dietzel, S., Mechtler, K., Durbin, R., Stewart, A. F., Peters, J. M., Buchholz, F. & Hyman, A. A. 2008. BAC TransgeneOmics: a high-throughput method for exploration of protein function in mammals. Nat Methods, 5, 409–15.

57. Ran, F. A., Hsu, P. D., Wright, J., Agarwala, V., Scott, D. A. & Zhang, F. 2013. Genome engineering using the CRISPR-Cas9 system. Nat Protoc, 8, 2281–2308.

58. Ray, N., Kunkle Br, Hamilton-Nelson, K., Kurup, J., Rajabli, F., Cosacak, M., Kizil, C., Jean-Francois, M., Cuccaro, M., Reyes-Dumeyer, D., Cantwell, L., Kuzma, A., Vance, J. M., Gao, S., Hendrie, H., Baiyewu, O., Ogunniyi, O., Akinyemi, R., ALZHEIMER’S DISEASE GENETICS Consortium, Lee, W.-P., Martin, E., Wang, L.-S., Beecham, G., Bush, W., Farrer, L., Haines, J., Byrd, G., Schellenberg, G., Mayeux, R., PERICAK-Vance, M. & Reitz, C. 2023. Extended genome-wide association study employing the African genome resources panel identifies novel susceptibility loci for Alzheimer’s disease in individuals of African ancestry. Alzheimers Dement.

59. Rodrigues, V. L., Dolde, U., Sun, B., Blaakmeer, A., Straub, D., Eguen, T., Botterweg-Paredes, E., Hong, S., Graeff, M., Li, M. W., Gendron, J. M. & Wenkel, S. 2021. A microProtein repressor complex in the shoot meristem controls the transition to flowering. Plant Physiol, 187, 187–202.

60. Siddiqui, T., Bhattarai, P., Popova, S., Cosacak, M. I., Sariya, S., Zhang, Y., Mayeux, R., Tosto, G. & Kizil, C. 2021. KYNA/Ahr Signaling Suppresses Neural Stem Cell Plasticity and Neurogenesis in Adult Zebrafish Model of Alzheimer’s Disease. Cells, 10.

61. Silva, J. C., Gorenstein, M. V., Li, G. Z., Vissers, J. P. & Geromanos, S. J. 2006. Absolute quantification of proteins by LCMSE: a virtue of parallel MS acquisition. Mol Cell Proteomics, 5, 144–56.

62. Staudt, A. C. & Wenkel, S. 2011. Regulation of protein function by ’microProteins’. EMBO Rep, 12, 35–42.

63. Stein, C. S., Jadiya, P., Zhang, X., Mclendon, J. M., Abouassaly, G. M., Witmer, N. H., Anderson, E. J., Elrod, J. W. & Boudreau, R. L. 2018. Mitoregulin: A lncRNA-Encoded Microprotein that Supports Mitochondrial Supercomplexes and Respiratory Efficiency. Cell Rep, 23, 3710–3720 e8.

64. Tayran, H., Yilmaz, E., Bhattarai, P., Min, Y., Wang, X., Ma, Y., Wang, N., Jeong, I., Nelson, N., Kassara, N., Cosacak, M. I., Dogru, R. M., Reyes-Dumeyer, D., Stenersen, J., Reddy, J., Qiao, M., Flaherty, D., Gunasekaran, T. I., Yang, Z., Jurish-Yaksi, N., Teich, A. F., Kanekiyo, T., Tosto, G., Vardarajan, B., Is, O., Ertekin-Taner, N., Mayeux, R. & Kizil, C. 2024. ABCA7-dependent Neuropeptide-Y signalling is a resilience mechanism required for synaptic integrity in Alzheimer’s disease. Cell Genomics, 4, 100642.

65. Turgutalp, B., Bhattarai, P., Ercetin, T., Luise, C., Reis, R., Gurdal, E., Isaak, A., Biriken, D., Dinter, E., Sipahi, H., Schepmann, D., Junker, A., Wünsch, B., Sippl, W., Gulcan, H., Yarim, M. & Kizil, C. 2022. Discovery of Potent Cholinesterase Inhibition-Based Multi-Target-Directed Lead Compounds for Synaptoprotection in Alzheimer’s Disease. J Med Chem.

66. Turgutalp, B. & Kizil, C. 2024. Multi-target drugs for Alzheimer’s disease. Trends Pharmacol Sci, 45, 628–638.

67. Wang, L., Fan, J., Han, L., Qi, H., Wang, Y., Wang, H., Chen, S., Du, L., Li, S., Zhang, Y., Tang, W., Ge, G., Pan, W., Hu, P. & Cheng, H. 2020. The micropeptide LEMP plays an evolutionarily conserved role in myogenesis. Cell Death Dis, 11, 357.

68. Wu, Q., Zhong, S. & Shi, H. 2022. MicroProteins: Dynamic and accurate regulation of protein activity. J Integr Plant Biol, 64, 812–820.

69. Xu, H., Somers, Z. B., Robinson, M. L., 2ND & Hebert, M. D. 2005. Tim50a, a nuclear isoform of the mitochondrial Tim50, interacts with proteins involved in snRNP biogenesis. BMC Cell Biol, 6, 29.

70. Xu, W., Deng, B., Lin, P., Liu, C., Li, B., Huang, Q., Zhou, H., Yang, J. & Qu, L. 2020. Ribosome profiling analysis identified a KRAS-interacting microprotein that represses oncogenic signaling in hepatocellular carcinoma cells. Sci China Life Sci, 63, 529–542.

71. Yang, W., Zou, Y., Zhang, M., Zhao, N., Tian, Q., Gu, M., Liu, W., Shi, R., Lu, Y. & Yu, W. 2015. Mitochondrial Sirt3 Expression is Decreased in APP/PS1 Double Transgenic Mouse Model of Alzheimer’s Disease. Neurochem Res, 40, 1576–82.

72. Zhang, S., Reljic, B., Liang, C., Kerouanton, B., Francisco, J. C., Peh, J. H., Mary, C., Jagannathan, N. S., Olexiouk, V., Tang, C., Fidelito, G., Nama, S., Cheng, R. K., Wee, C. L., Wang, L. C., DUEK Roggli, P., Sampath, P., Lane, L., Petretto, E., Sobota, R. M., Jesuthasan, S., TUCKER-Kellogg, L., Reversade, B., Menschaert, G., Sun, L., Stroud, D. A. & Ho, L. 2020. Mitochondrial peptide BRAWNIN is essential for vertebrate respiratory complex III assembly. Nat Commun, 11, 1312.

73. Zhang, W., Gu, G.-J., Shen, X., Zhang, Q., Wang, G.-M. & Wang, P.-J. 2015. Neural stem cell transplantation enhances mitochondrial biogenesis in a transgenic mouse model of Alzheimer’s disease-like pathology. Neurobiology of Aging, 36, 1282–1292.

74. Zhao, Z., Bo, Z., Gong, W. & Guo, Y. 2020. Inhibitor of Differentiation 1 (Id1) in Cancer and Cancer Therapy. Int J Med Sci, 17, 995–1005.

